# Hierarchy, Morphology, and Adaptive Radiation: a Test of Osborn’s Law in the Carnivora

**DOI:** 10.1101/285700

**Authors:** Graham J. Slater, Anthony R. Friscia

## Abstract

Henry Fairfield Osborn’s law of adaptive radiation was intended to explain the early proliferation of morphological and functional variation in diversifying clades. Yet, despite much theoretical development and empirical testing, questions remain regarding the taxonomic levels at which adaptive radiation occurs, the traits involved, and its frequency across the tree of life. Here, we evaluate support for this “early burst” model of adaptive radiation in 14 ecomorphological traits plus body mass for the extant mammalian order Carnivora. Strong support for an early burst adaptive radiation is recovered for molar grinding area, a key proxy for diet. However, we find no evidence for early burst–like dynamics in body mass or multivariate trait data, suggesting a decoupling of evolutionary modes among traits driven by dietary specialization. Furthermore, the signal of an early burst is only recovered for Carnivora, and not in family–level clades. The lack of support for the early burst model of morphological adaptive radiation in previous phylogenetic studies may be a consequence of focusing on the wrong traits at the wrong taxonomic levels. Osborn’s law predicted that adaptive radiation should be hierarchically structured, and the search for its signature and understanding of its prevalence will require a renewed focus on functional traits and their evolution over higher-level clades.

## Introduction

Since the turn of the twentieth century, adaptive radiation has been a dominant paradigm for understanding the uneven distribution of morphological diversity across the tree of life, and in clades ranging from arthropods (Gould, 1989; Briggs et al., 1992) to lungfishes (Westoll, 1949) and island lizards (Losos, 2009). Arising initially from Henry Fairfield Osborn’s “Law of Adaptive Radiation” (Osborn, 1902), and undergoing theoretical development by paleobiologist (Simpson, 1944, 1953; Valentine, 1969, 1980; Walker and Valentine, 1984; Gould, 1989), quantitative geneticists (Lande, 1976; Arnold et al., 2001), and macroevolutionary biologists (Schluter, 1996b, 2000; Gavrilets and Vose, 2005; Harmon et al., 2010), this model posits that increased ecological opportunity, resulting from extinction of competitors, colonization of a novel environment, or the evolution of a key innovation, triggers rapid expansion of a diversifying clade across the adaptive landscape and, in turn, morphospace (Gavrilets and Vose, 2005; Gavrilets and Losos, 2009; Losos and Mahler, 2010; Yoder et al., 2010). As niches fill and ecological opportunity is exhausted, the rate at which the clade spreads across the adaptive landscape should slow, yielding a characteristic signal that has come be known as an “Early Burst” of morphological evolution.

Robust empirical support for the early burst hypothesis is conspicuously lacking however. Disparity, measured as per-interval morphological variance, reaches a peak early in the fossil records of many clades (e.g., Knoll et al., 1984; Foote, 1992, 1994, 1995; Lupia, 1999; Boyce, 2005; Ruta et al., 2006; Hughes et al., 2013), but this pattern could equally arise as the result of constraints that restrict a clade to repeatedly revisit the same limited set of morphologies (Foote, 1996; Gavrilets, 1999). These very different evolutionary modes – declining evolutionary rates versus constant rates in a bounded space – are distinguishable if the phylogenetic relationships among taxa are known, as true early bursts should leave a strong phylogenetic signal in the distribution of trait values, while limits to disparity should result in homogenization of variation within and among clades, in turn reducing phylogenetic signal (Revell et al., 2008; Slater, 2015b). But, while early bursts of morphological diversification have been recovered in some clades using phylogenetic methods (e.g., Cooper and Purvis, 2010; Slater et al., 2010; Benson et al., 2014; Derryberry et al., 2011; Astudillo-Clavijo et al., 2015; Price et al., 2016), they are far from common and an analysis of 44 extant clades, including some of the most iconic examples of adaptive radiation, revealed a dearth of support for early burst dynamics in both size and shape data (Harmon et al., 2010). This has led some authors to question whether the early burst model of adaptive radiation is truly as widespread as suggested by the fossil record, and to a view that morphological evolution is essentially unbounded in space and time (Venditti et al., 2011). As a result, the study of tempo and mode in morphological evolution, at least from a phylogenetic perspective, has begun to swing away from the goal of identifying law-like generalities about how evolution works (Gould, 1980; Marshall, 2014) and towards more *ad hoc* objectives such as identifying rate shifts (Eastman et al., 2011; Venditti et al., 2011; Rabosky, 2014) or the presence of adaptive peaks (Ingram and Mahler, 2013; Mahler et al., 2013; Uyeda and Harmon, 2014; Khabbazian et al., 2016) that are specific to individual clades.

Before discarding the early burst paradigm, it is worth reconsidering its origins. Osborn’s original law of adaptive radiation (Osborn, 1902) was ostensibly a pair of ideas about how ecomorphological diversification proceeds at different levels of the phylogenetic hierarchy. In his framework, ecological opportunity drove a burst of diversification in locomotor and feeding morphologies early in clade history, consistent with the early burst paradigm, that resulted in a *general* adaptive radiation. But, Osborn also noted that subsequent competition among contemporaneous taxa occupying similar adaptive zones would drive the continuous evolution of characteristics that facilitate resource partitioning, resulting in ongoing *local* adaptive radiations. Paleobiologists since Osborn’s time have developed conceptually similar models to understand how rates or modes of morphological evolution vary through clade history (e.g., Gould, 1989; Simpson, 1944, 1953; Valentine, 1980) but the unifying concepts, often missed in phylogenetic comparative studies (but see Humphreys and Barraclough, 2014), are that hierarchy and traits matter (Jablonski, 2007; Erwin, 2000); while higher level clades tend to represent patterns of morphological and ecological diversification that are consistent with general adaptive radiation, lower level clades, such as vertebrate families and genera, almost always represent local adaptive radiations, a “filling–in” of already occupied ecospace, and their diversity is rarely predictive of morphological disparity in general terms (Foote, 1993, 1995, 1996, 1997; Valentine, 1980, 1969). Thus, although Schluter’s (2000) ecological theory of adaptive radiation emphasizes the importance of ecological opportunity in driving adaptive radiation, it focuses on too low a level for early bursts of morphological evolution to be an expected outcome and it comes as little surprise that this pattern is rare among low-level phylogenetic datasets examined to date, as in Harmon et al. (2010). Given the need for a phylogenetic context when examining patterns of morphological evolution (Foote 1996) and the wide availability of suitable methods for detecting early bursts (Blomberg et al., 2003; Freckleton and Harvey, 2006; Harmon et al., 2003, 2010), the more pressing question is not whether early bursts occur, but rather at what level and in which traits they are found, and then under what circumstances.

Appropriate systems for testing Osborn’s hierarchical model of adaptive radiation should meet three distinct criteria. First, they should be higher level clades, such as classes or orders, that encompass several lower level clades. Second, they should possess a well–resolved phylogeny with branch lengths, and third, they should exhibit a range of ecomorphological variation that can be both quantified and directly related to resource use. We here study ecomorphological macroevolutionary dynamics using terrestrial representatives of the mammalian order Carnivora as just such a model system. Carnivora is diverse, comprising over 200 species in 13 extant families (exclusive of the aquatic pinnipeds) and its phylogenetic relationships have been extensively studied (Gaubert and Veron, 2003; Bardeleben et al., 2005; Flynn et al., 2005; Gaubert and Cordeiro-Estrela, 2006; Fulton and Strobeck, 2006, 2007; Johnson et al., 2006; Koepfli et al., 2006, 2007, 2008; Krause et al., 2008; Patou et al., 2009; Eizirik et al., 2010). Furthermore, the broad ecological diversity in dietary habits among carnivorans (Ewer, 1973), can be directly related to a rich suite of ecomorphological traits that can readily measured and that have demonstrable form-function relationships (Van Valkenburgh, 1988, 2007; Van Valkenburgh and Koepfli, 1993; Friscia et al., 2007). Here, we use a novel species–level phylogeny and ecomorphological dataset consisting of craniodental functional indices and body mass to ask which, if any, ecomorphological traits show signals consistent with an early burst of morphological evolution and at what phylogenetic levels these signals emerge. We find that Osborn’s model of general adaptive radiation is supported at the level of Carnivora for traits that have direct links to the broad dietary niche (i.e., carnivory, omnivory, herbivory) occupied by constituent species, but not for traits related to size. However, the signal of a general adaptive radiation is not recovered at lower levels for any trait, suggesting that the early burst pattern may be an emergent property of complex within-clade dynamics rather than an actual process. By modelling the evolution of the evolutionary rate matrix describing evolutionary correlations between molar grinding area and body mass, we also show that axes of local adaptive radiation may differ among sub–clades of a general adaptive radiation.

## Materials and methods

### Phylogeny Reconstruction

We inferred a new, time-scaled phylogeny for 235 extant and recently extinct terrestrial carnivorans, omitting pinnipeds (seals, sealions and walrus) as our trait evolutionary analyses focus on terrestrial clades. Full details of phylogenetic methods are provided in the Supplementary Methods but, briefly, we downloaded 30 thirty nuclear and all available protein coding mitochondrial loci from Genbank and jointly estimated phylogenetic topology and branch lengths in a Bayesian framework as implemented in BEAST v2.3.2 (Bouckaert et al., 2014) under a relaxed log-normal molecular clock (Drummond et al., 2006) and using the Fossilized Birth–Death (FBD) model as a tree prior (Heath et al., 2014; Gavryushkina et al., 2014, 2016). Following the approach described in Law et al. (2017) for musteloid carnivorans, we included 250 fossil carnivorans as terminal taxa in our matrix with all characters coded as “?” and occurrence times specified as uniform age range corresponding to described stratigraphic ranges. To reduce treespace for the topology search while avoiding the need to make arbitrary decisions regarding relationships of fossil taxa, we specified a minimal set of topology constraints based on the most inclusive placements possible as ascertained from published phylogenetic analyses. We ran three independent chains, each of 100 million generations, sampling every 10,000 generations. After visually checking for convergence of likelihoods and parameters using the TreeAnnotator software, we discarded a chain-specific appropriate number of samples as burn-in and combined the posterior samples of trees before pruning fossil taxa and producing a Maximum Clade Credibility (MCC) tree from the retained posterior sample. This tree was used for subsequent macroevolutionary analyses.

### Morphological Data Collection

Our trait evolution analyses are based on 14 functional variables plus body mass (Figure 1, Table 1). The ecomorphological traits that we use describe aspects of craniodental morphology associated with food acquisition and processing that have been shown to be important discriminators of diet and predatory behavior in previous studies (Friscia et al., 2007; Meachen-Samuels and Van Valkenburgh, 2009; Sacco and Van Valkenburgh, 2004; Van Valkenburgh, 1988, 1991; Van Valkenburgh and Koepfli, 1993). The dataset is based on measurements from 1583 museum specimens representing 198 terrestrial carnivoran species (*n* = 8, range = 1-25). Measurements for a subset of specimens (*n* = 425) were taken from a previous study of small carnivoran ecomorphology (Friscia et al., 2007) while the remaining measurements were collected specifically for this project using Mitutuyo (c) digital calipers to 0.01mm precision. We excluded the aardwolf (*Proteles cristatus*) due to its peg-like teeth that cannot be measured for our ecomorphological traits, and the bat-eared fox (*Otocyon megalotis*) due to its duplicated upper first and lower second molars. Where possible, only wild-caught specimens were measured due to well-described skeletal abnormalities associated with captive carnivorans (O’Regan and Kitchener, 2005; Hartstone-Rose et al., 2014), though dental measurements were taken from some zoo animals to increase sample sizes. We obtained body masses, in kilograms, for all available terrestrial carnivoran species in the panTHERIA database (Jones et al., 2009) and supplemented these with data from the literature for missing species. We converted our mass estimates to a linear scale by taking their cube roots and subsequently natural log-transformed them for analysis.

**Figure 1:**
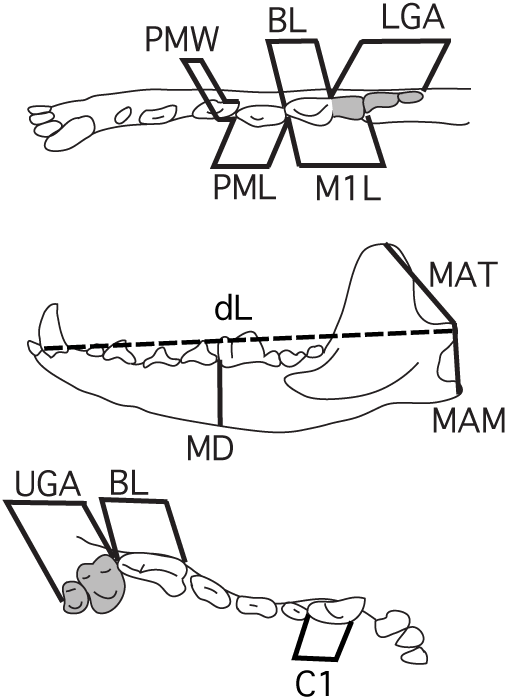
Measurements used to generate ecomorphological metrics. Top: occlusal view of lower dentition. Middle: lingual view of lower dentition. Bottom: occlusal view of upper dentition. PMW, lower fourth premolar width; PML, lower fourth premolar length; BL, blade length of carnassial; M1L, length of m1; LGA, lower grinding area (occlusal area of talonid of m1 and post-carnassial dentition); MAM, moment arm of masseter muscle; MAT, moment arm of temporalis muscle; dL, dentary length; MD, mandibular depth; UGA upper grinding area (occlusal area of upper molars). Modified from Friscia et al.(2007).

**Table 1:** Functional traits, their meanings, and constituent measurements.

| Abbreviation | Meaning |
| --- | --- |
| C1 | Cross sectional shape of the upper canine, measured as width divided by anteroposterior length |
| P4S | Lower fourth premolar shape, measured as maximum width of p4 divided by its maximum length |
| P4Z | Relative length of fourth lower premolar, measured as the maximum length of p4 divided by dentary length (measured as in M1BS) |
| P4P | Relative size of the protocone of the upper fourth premolar (carnassial), measured as the ratio of maximum width of P4 divided by maximum length of P4 |
| RBL | Relative blade length of lower first molar (m1 carnassial), measured as the ratio of trigonid length to total anteroposterior length of m1 |
| M1BS | m1 blade size relative to dentary length, measured as the length of the trigonid of m1 (carnassial) divided by dentary length. |
| RLGA | Relative lower grinding area, measured as the square root of the summed areas of the m1 talonid and m2 (if present) divided by the length of the m1 trigonid. Area was estimated as the product of maximum width and length of the talonid of m1 and m2, respectively |
| M2S | m2 size relative to dentary length, measured as the square root of m2 area (if present) divided by dentary length. Tooth area measured as in RLGA and dentary length (measured as in M1BS). If no m2 was present in the taxon, M2S was recorded as zero |
| RUGA | Relative upper grinding area, measured as the square root of the summed areas of M1 and M2 (if present) divided by the anteroposterior length of P4 (upper carnassial). Area was estimated by the product of width and length of M1 and M2, respectively |
| UM21 | Square root of upper second molar area (if present) divided by square root of upper first molar area. Areas estimated as RUGA. If no M2 was present in a taxon, UM21 was recorded as zero |
| MAT | Mechanical advantage of the temporalis muscle, measured as the distance from the mandibular condyle to the apex of the coronoid process divided by dentary length (measured as in M1BS) |
| MAM | Mechanical advantage of the masseter muscle, measured as the distance from the mandibular condyle to the ventral border of the mandibular angle divided by dentary length (measured as in M1BS) |
| IxP4 | Second moment of area of the dentary at the interdental gap between the third and fourth lower premolars relative to dentary length. Used as an estimate of resistance of the dentary to bending. Second moment of area was calculated using the formula $I_x = (\pi D_x D_y^3)/64$, where $D_x$ is maximum dentary width and $D_y$ is maximum dentary height at the p3-p4 interdental gap. $I_x$ relative to dentary length was then estimated as the fourth root of $I_x$ divided by dentary length (measured as in M1BS) |
| IxM2 | same as IxP4 but at the interdental gape between the first and second lower molar or, if m2 is absent, immediately posterior to m1. |
| $\ln(\sqrt[3]{mass})$ | the natural logarithm of the cube root of mass, in grams |

### Modeling Ecomorphological Evolution

We fit three standard models of morphological evolution to the species means for each of our 15 traits (14 ecomorphological traits plus body size) using the fitContinuous() function in the geiger v.2.0.6 library (Pennell et al., 2014). To understand whether and how hierarchy affects inference of evolutionary mode, we fit models to the entire carnivoran dataset, and to family-level clades containing *>* 5 species.

Brownian Motion (BM) is often invoked to describe a trait evolving by drift (i.e., as a random walk at a constant rate) but can also describe processes in which evolution tracks a fluctuating or rapidly shifting adaptive peak (Revell et al., 2008). We here interpret BM as any process for which trait covariances among taxa are well predicted by their phylogenetic covariances. A Single Stationary Peak (SSP) model is a single peak Ornstein-Uhlenbeck (OU) process where the root state and trait attractor are equivalent (Hansen, 1997). This model can describe trait evolution under attraction to a single adaptive peak but may also fit well to datasets generated under constrained or bounded evolution. The early burst (EB) model (Blomberg et al., 2003; Harmon et al., 2010) is a time-heterogeneous rate model of morphological evolution where rate decays exponentially through time from its initial value, resulting in a strong phylogenetic partitioning of morphological variation. As a result, this model provides a good approximation of Osbornian general adaptive radiation. Measurement error was accounted for in univariate analyses by adding sampling variance to the diagonals of the variance-covariance matrix. Because sample sizes were small for some species, we computed a pooled variance for each trait and then computed the sampling variance for the *i*th species as the pooled variance divided by its sample size, *n_i_*. Because we did not have within-species estimates of sampling error for body mass, we assigned each species a standard error of 0.0345, following Harmon et al. (2010). Relative model fit was assessed by computing small-sample corrected Akaike Weights, *w_A_*, for each model.

Because morphology is inherently multivariate, it is possible that adaptive radiations occur in multivariate ecomorphospace, rather than along individual trait axes. Under multivariate Brownian motion with *k* traits, we must estimate a vector of *k* phylogenetic means, or root states, and a *k* x *k* matrix, **R**, the diagonal elements of which give the evolutionary rates of each trait and off-diagonal elements give the evolutionary covariances that describe how traits change in relation to one another. For this reason, **R** is often referred to as the evolutionary rate matrix (a macroevolutionary equivalent of the **P**-matrix from quantitative genetics). We fit multivariate versions of the three models described above to the complete, 15 trait dataset using the fast method proposed by Freckleton (2012), which uses phylogenetically independent contrasts (Felsenstein, 1985) in place of raw trait data and thus avoids the need to repeatedly invert the outer product of the phylogenetic covariance matrix and evolutionary rate matrix. Using this approach, it is straightforward to fit multivariate versions of the SSP or EB model by appropriately transforming the branch lengths of the phylogeny prior to computing the contrasts variances (Freckleton, 2012), provided that the tree is ultrametric (Slater, 2014). It should be noted that our non-BM multivariate models assume that the effect of the evolutionary model’s parameters act uniformly across traits, for example that the evolutionary rate of each trait decays at the same rate under a multivariate EB model.

### Macroevolutionary Covariance of Size and Ecomorphology

We used a Bayesian approach, as implemented in the R library ratematrix (Caetano and Harmon, 2017a,b), to understand whether the rate matrix itself evolves in response to shifts in dietary ecology and whether this can explain different macroevolutionary signals in different traits. We used an evolutionary model-based approach to classify carnivorans to dietary regimes, rather than the more traditionally approach of using literature review. This is because diet can vary considerably and continuously both within and among species, with the net effect that dietarily dissimilar taxa become lumped into a common but meaningless omnivore regime (Davis and Pineda-Munoz, 2016; Pineda-Munoz and Alroy, 2014; Pineda-Munoz et al., 2017). Using Relative Lower Grinding Area as a focal trait due to its well–demonstrated link to dietary resource use (Van Valkenburgh, 1988, 1991), we inferred the presence and location of dietary regimes under an Ornstein-Uhlenbeck process as implemented in the R library l1ou (Khabbazian et al., 2016). This approach uses a phylogenetic LASSO (Least Absolute Shrinkage and Selection Operator) to identify an optimal set of evolutionary regimes on a phylogenetic tree, given a set of trait values for the terminal taxa. In this respect it is similar to the SURFACE approach of Ingram and Mahler (2013), which uses a step-wise AIC procedure to identify unique evolutionary regimes under an OU process, but is substantially faster for larger datasets. We performed an initial analysis to identify distinct evolutionary regimes with a maximum of 20 shifts enforced and using the Bayesian Informaiton Criterion for model selection. We then performed a second analysis on the resulting model output to identify convergent regimes that could be collapsed into one another. The final set of regimes can be thought of as representing large shifts in morphology relative to background evolutionary rates that, in turn, correspond to shifts in diet and we consider them analogous to Simpson’s (1944) notion of major adaptive zones.

To sample evolutionary rate matrices for each dietary regime, we first transformed each vector of trait values to a 0-1 scale and placed broad, gaussian priors on the root states of each trait, with a mean corresponding to the arithmetic mean of the trait and a standard deviation of 1.5 times the empirical standard deviation. For computational efficiency we restricted our analyses here to sample the 2X2 **R** matrix describing evolutionary correlations between Relative Lower Grinding Area and 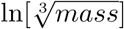. This choice is motivated by the fact that a large number of macroevolutionary analyses have focused on body mass as a proxy for ecomorphology when attempting to understand adaptive radiation. We set a lognormal prior (ln[mean] = −2.3, sd = 1) on the standard deviations (square roots of diagonal elements) of the evolutionary rate matrix and a flat prior on the correlation matrices. We ran two Markov chains for 1 million generations each and checked for convergence by ensuring potential scale reductions factors reached 1 and that effective sample sizes for all parameters were greater than 200. We then discarded the first 20% of each run as burn-in and thinned the posterior by retaining every 1000th sample.

The use of a Bayesian approach to sample from the joint posterior distribution allows us to make use of Bayesian tools for comparing rate matrices among different evolutionary regimes. We first used Ovaskainen et al.’s (2008) method for assessing the difference between joint posterior distributions of covariance matrices for two groups. Because a covariance matrix defines a multivariate normal distribution, we can assess the distance *d*(**A**, **B**) between two matrices by simulating many datasets from **A** and computing the proportion of cases in which these simulated data have a higher probability of originating from **B**. To assess statistical support for the difference between two posterior samples, pairs of covariance matrices are repeatedly drawn from the joint posterior distributions of regimes A and B. We first measure the distances between pairs of matrices from the same regime and subtract from their sum the summed distance between joint samples from the two regimes,

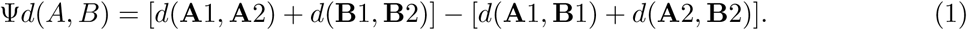

If inter-regime variation (the second term on the right side of equation 1) is large relative to within-regime variation (the first term), that is the difference between regimes is substantial even after accounting for uncertainty in within–regime estimates, then Ψ*d*(*A, B*) will be negative and a statistical test of whether the two regimes share a common **R** can be computed as *P* [Ψ*d*(*A, B*) *>* 0]. Because this test is one-tailed, we accept the two samples as representing different distributions if *P* [Ψ*d*(*A, B*) *>* 0] *<* 0.05

Ovaskainen et al.’s test tells us whether **R** differs among regimes but not whether it differs in shape, size, or orientation. Matrix size is determined by the diagonal elements of **R**, which give the evolutionary rates for each trait. The sum of the diagonals, tr(**R**), therefore gives the rate of dispersion in multivariate space (Haber, 2016). The shape, or eccentricity, of the clade’s dispersion in multivariate space can be characterized by the relative standard deviation of the eigenvalues (rSDE) of **R** (Van Valen, 1974; Haber, 2016, see also Kirkpatrick 2009). Values of rSDE close to zero indicate that variance is evenly spread over the eigenvectors of **R**, meaning that constraints due to evolutionary covariation are weak and that the clade is free to evolve in any direction in morphospace. Values of rSDE approaching 1 indicate that covariation among traits is strong and that that the clade is constrained to evolve along a single axis. We computed, for each pairwise comparison among dietary regimes, the posterior probability that the rate of dispersion and rSDE were greater in regime A than for regime B as the proportion of cases in which tr(**R**) and rSDE of samples from the joint posterior of regime A were larger than those of regime B. Finally, we determined whether **R** differs in orientation among dietary regimes by calculating the angle between principal eigenvectors from the joint posterior sample. We used Equation 1 to determine whether the inter-regime difference in the orientation of **R** is greater than the uncertainty associated with each within-regime posterior sample. The probability that the regimes share a common orientation is then given as *P* [Ψ∠(*A, B*) *>* 0].

## Results

### Phylogeny Reconstruction

The topology of the maximum clade credibility tree from our BEAST analysis (Figure 2) is generally well-supported, with 94% nodes having posterior probabilities greater than 0.9, and is congruent with previous analyses of carnivoran phylogeny. Uncertainty in divergence time estimates was low (mean 95% Highest Posterior Density (HPD) interval = 2.56 Myr) but increased for older nodes, and was also elevated in areas of the tree that lack phylogenetically corroborated fossil taxa (e.g., the families Viverridae, Herpestidae, and Eupleridae) and/or areas with sparse molecular coverage such as the genera *Prionodon* and *Mydaus*. We infer an Early Eocene origin for crown Carnivora (mean age = 48.2 Ma, 95% HPD interval 45.1-52.6 Ma), with crown Caniformia originating shortly after (mean age = 46.1Ma, 95% HPD = 42.3 - 49.6 Ma). Crown Feliformia originated closer to the Eocene/Oligocene boundary (32.8Ma, 95% HPD = 29.1-37 Ma), and most extant families began to diversify during the mid to late Miocene or Pliocene. Only Procyonidae and Mephitidae have 95% HPD intervals that extend into the Oligocene.

**Figure 2:**
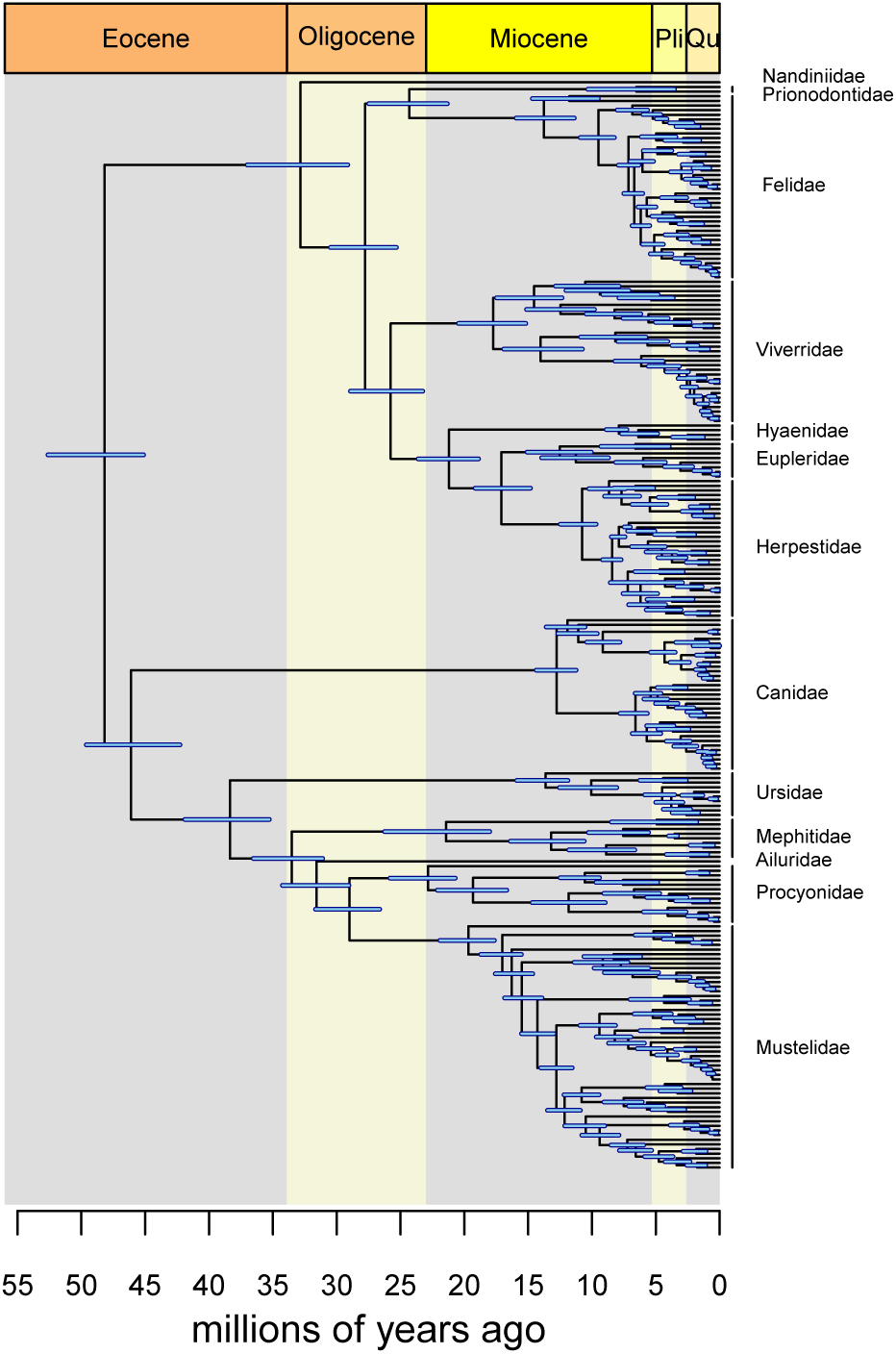
Maximum clade credibility tree of extant and recently extinct carnivorans inferred under a relaxed molecular clock and fossilized birth-death tree prior. Blue bars indicate 95% HPD intervals

### Modeling Ecomorphological Evolution

Early burst is the best fitting evolutionary model for a subset of dental traits associated with food-processing (Figure 3; Table 2), with strongest support (*w_A_ >* 0.95) recovered in grinding-related traits such as relative lower (RLGA), and upper (RUGA) grinding areas, and relative size of the upper second molar (UM21). Maximum Likelihood Estimates (MLEs) for the rate half-lives, that is the time for evolutionary rate to halve its initial value (Slater and Pennell, 2014), suggest moderate to slowly declining rates (*t*_1*/*2_ = 4.2 - 39.8 Myr). The single stationary peak model was recovered as best fitting model for several traits, but only those associated with bite strength (e.g., MAT, MAM), and canine tooth shape yielded moderate to strong support (*w_A_ >* 0.85). MLEs for the *α* parameter in these SSP models were small, corresponding to phylogenetic half-lives, a measure of the influence of the optimum relative to diffusion (Hansen, 1997), of between 10 and 35 Myr. Support for this model can therefore be interpreted as indicating multiple peak shifts over carnivoran history that have reduced phylogenetic signal in trait values, rather than strong constraint that prevented trait diversification. Brownian motion, though the best fitting model for premolar shape, lower carnassial blade size and body mass, was never strongly supported over other models.

**Figure 3:**
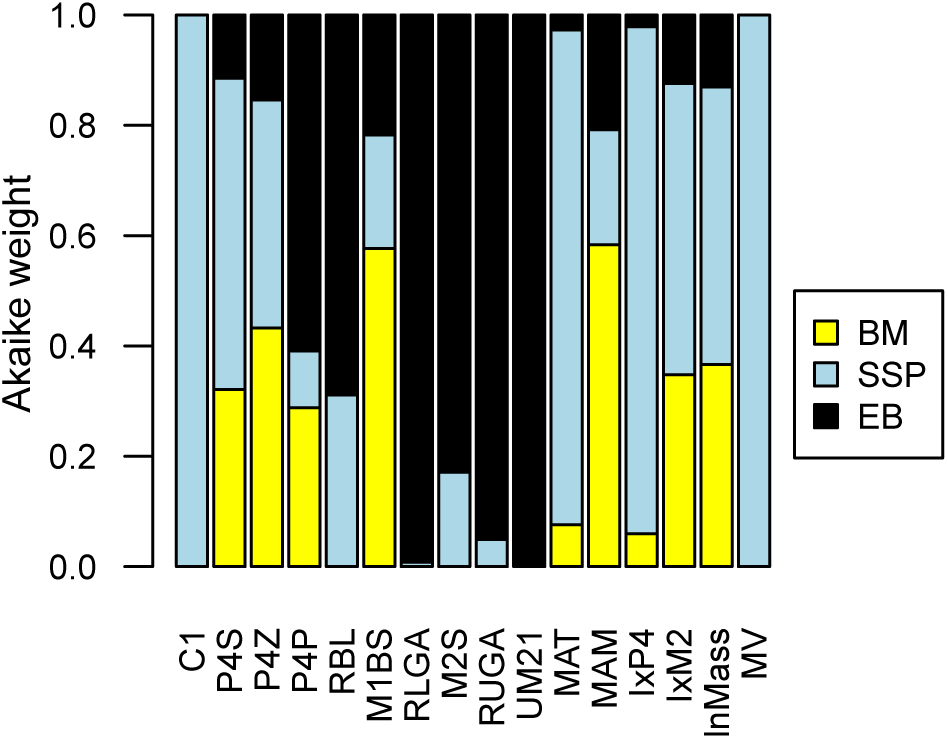
Stacked barplots of Akaike Weights for the three models of morphological evolution (BM, Brownian motion; SSP, single stationary peak; EB, early burst) fitted to each of the 14 ecomorphological traits, body mass, and the multivariate (MV) dataset on the maximum clade credibility tree. The relative height of the color bars indicate relative support for the corresponding model given the trait

**Table 2:** Best fitting models for each ecomorphological trait fitted to the entire Carnivora, along with associated parameter estimates and relative model support. *σ*^2^ and *θ* for MV data are vectors, rather than single values, and so are omitted here.

| trait | best model | $\alpha$ | $r$ | $\sigma^2$ | $\theta$ | $w_A$ |
| --- | --- | --- | --- | --- | --- | --- |
| C1 | SSP | 0.09 | - | $8.10 * 10^{-4}$ | 0.71 | 1.00 |
| P4S | SSP | 0.02 | - | $6.00 * 10^{-4}$ | 0.56 | 0.56 |
| P4Z | BM | - | - | $1.21 * 10^{-4}$ | 0.09 | 0.43 |
| P4P | EB | - | -0.04 | $3.96 * 10^{-4}$ | 0.71 | 0.61 |
| RBL | EB | - | -0.02 | $9.43 * 10^{-4}$ | 0.64 | 0.69 |
| M1BS | BM | - | - | $2.29 * 10^{-5}$ | 0.09 | 0.58 |
| RLGA | EB | - | -0.06 | $4.80 * 10^{-2}$ | 1.02 | 0.99 |
| M2S | EB | - | -0.03 | $5.79 * 10^{-5}$ | 0.06 | 0.83 |
| RUGA | EB | - | -0.04 | $1.48E * 10^{-2}$ | 1.04 | 0.95 |
| UM21 | EB | - | -0.08 | $4.48 * 10^{-2}$ | 0.51 | 1.00 |
| MAT | SSP | 0.03 | - | $1.78 * 10^{-4}$ | 0.27 | 0.90 |
| MAM | BM | - | - | $7.34 * 10^{-5}$ | 0.15 | 0.58 |
| IxP4 | SSP | 0.03 | - | $8.41 * 10^{-6}$ | 0.06 | 0.92 |
| IxM2 | SSP | 0.02 | - | $6.14 * 10^{-6}$ | 0.07 | 0.53 |
| $\ln(\sqrt[3]{mass})$ | BM | - | - | $1.09 * 10^{-2}$ | 0.56 | 0.45 |
| MV | SSP | 0.025 | - | - | - | >0.99 |

Strong support for the early burst mode of evolution is less conspicuous when analyses are conducted at lower phylogenetic levels (Figure 4). Among familes containing more than 5 taxa, an early burst of grinding area evolution is only recovered for Viverridae and support is weak at best (Figure 4i; *w_A_ >* 0.66). Well supported early bursts are recovered in a few clades, but for very different traits, for example relative size of the second molars (M2S, UM21) in Eupleridae (Figure 4b), mechanical advantage of the masseter muscle (MAM) in Procyonidae (Figure 4g), and relative length of the lower fourth premolar (PMZ) in Ursidae (Figure 4h).

**Figure 4:**
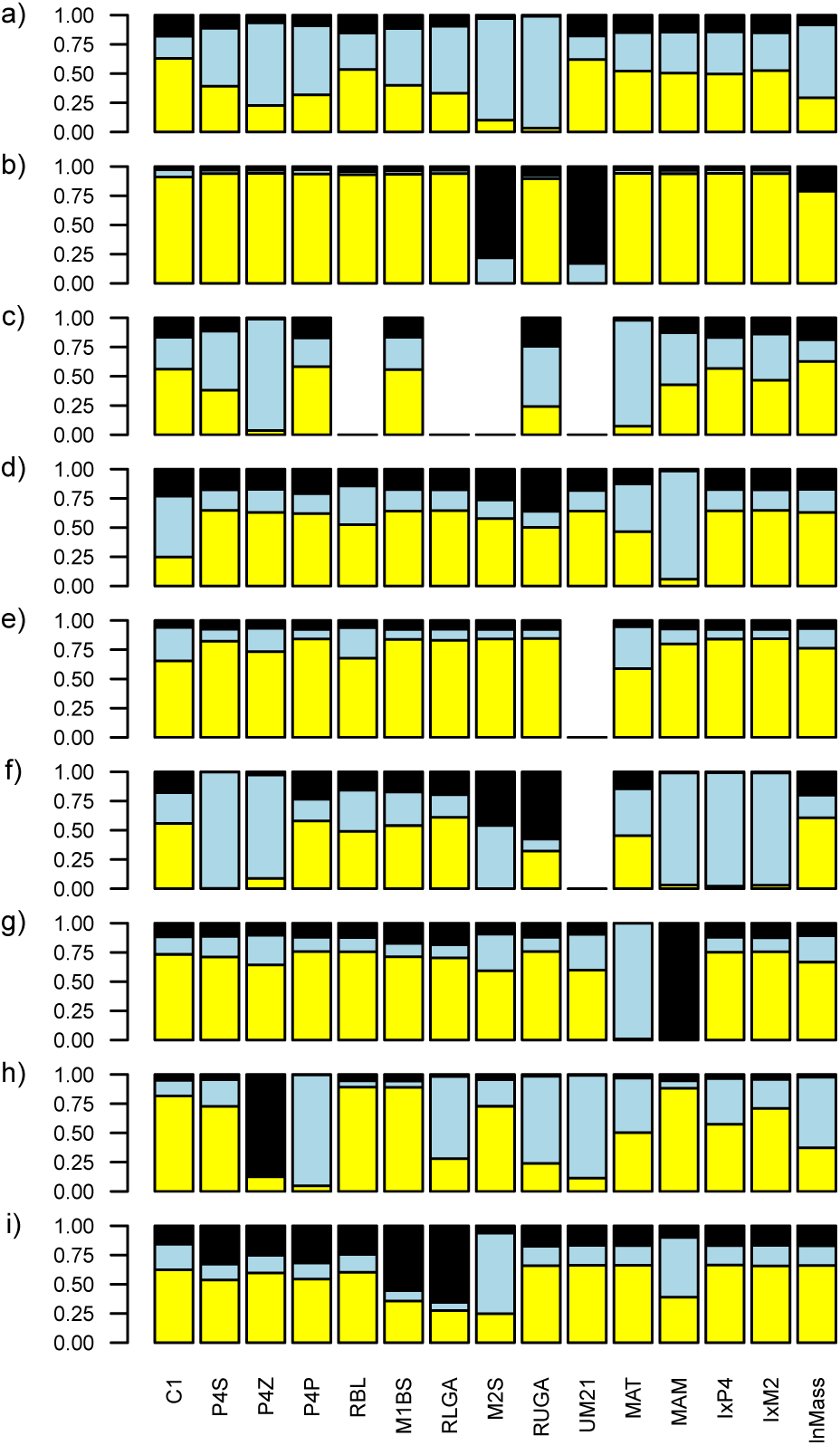
Stacked barplots of Akaike Weights for the three models of morphological evolution fitted to each of the 14 ecomorphological traits and body mass for each carnivoran family containing more than 5 species: a) Canidae, b) Eupleridae, c) Felidae, d) Herpestidae, e) Mephitidae, f) Mustelidae, g) Procyonidae, h) Ursidae, i) Viverridae. Colors correspond to those in Figure 3; Brownian motion: yellow; SSP: blue; Early burst: black.

We found no support for an early burst of multivariate morphological evolution, with an SSP model recovered as best-fitting. Visualization of the common carnivoran **R** matrix (Figure 5) reveals strong correlations in some traits (e.g, negative correlations between slicing and grinding function of the carnassial teeth) but most correlations are weak. Body mass, in particular, shows weak correlations with most ecomorphological traits.

**Figure 5:**
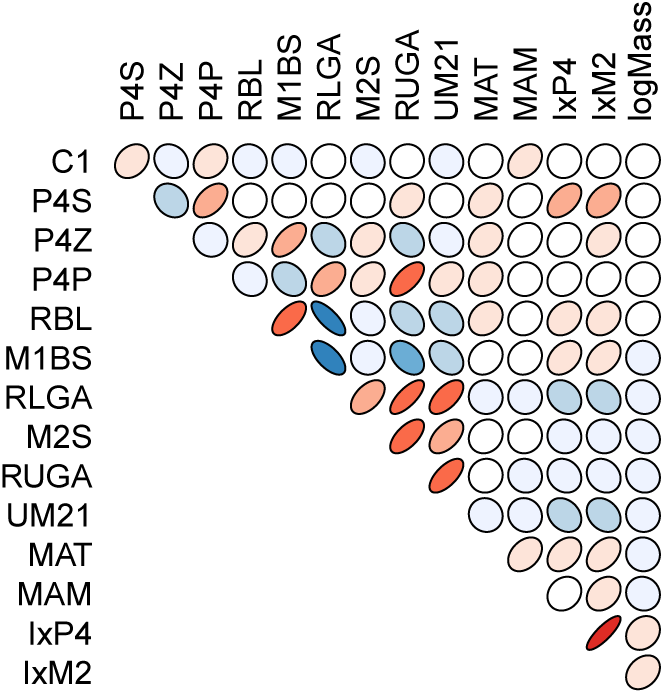
A plot of the correlation matrix derived from **R** for all carnivorans indicates strong correlations among functionally related traits but that the relationship between size and ecomorphology (final column or row) is weak. The shape of the ellipses indicates the strength of the correlation between the row and column traits, with more ellipical shapes indicating stronger correlations and circular shapes indicating weaker correlations. Hotter colors indicate stronger positive correlations while colder colors indicate stronger negative correlations

### Macroevolutionary Covariance of Size and Ecomorphology

We identified 12 independent dietary regime shifts in carnivorans using l1ou that were subsequently collapsed into one of five convergent regimes (Figure S1,2). These can be broadly summarized as (1) cat-like hypercarnivores (Felidae, Prionodontidae, Hyaenidae, *Cryptoprocta ferox*), 2) weasel-like hypercarnivores (most mustelid clades plus the herpestid genera *Paracynictis* and *Cynictis*), 3) omnivores and hard-object feeders (most arctoids, hemigaline viverrids, mungotine herpestids), 4) frugivores (Procyonidae exclusive of *Procyon* and *Bassariscus*), and 5) an ancestral regime that we recognize as mesocarnivorous (Canidae, most Viverridae, Eupleridae exclusive of *Cryptoprocta*, and remaining Herpestidae).

Bayesian joint estimation of **R** for the five dietary regimes resulted in good convergence and large effective sizes (*>>* 200) for all parameters except for regime 4 (frugivorous procyonids), which contained only 8 species; we therefore omit this group from subsequent pairwise comparisons. (Table 3). The 2×2 **R** matrices for the remaining dietary regimes can be well differentiated from one another based on Ovaskainen et al.’s (2008) distance test, though the comparison between omnivores and mesocarnivores is not significant at the *α* = 0.05 level (*p* = 0.09). Cat–like 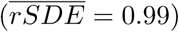 and weasel–like 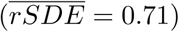 hypercarnivores have more eccentric **R**–matrices than omnivores 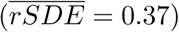 or mesocarnivores 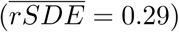, and cat–like hypercarnivores have more eccentric matrices than weasel–like taxa. Furthermore, the angles between the principal eigenvectors of the 2×2 **R**–matrices for ecologically distinct regimes differ significantly. The principal vectors of the two hypercarnivore regimes are almost perpendicular to those of omnivores (mean angles 75°and 68°, respectively) and are oriented in the direction of body mass variation (Figure 6). Comparisons between the principal vectors of hypercarnivore regimes (mean angle = 7°, p = 0.091) and between omnivores and mesocarnivores (mean angle = 22°, p = 0.272) are not significant. Taken together, these results indicate that hypercarnivores display limited ability to evolve in the direction of grinding area and disperse predominantly in the direction of size (Figure 6). Mesocarnivores and omnivores evolve dominantly in the direction of grinding area but retain the ability to evolve freely in the direction of size.

**Figure 6:**
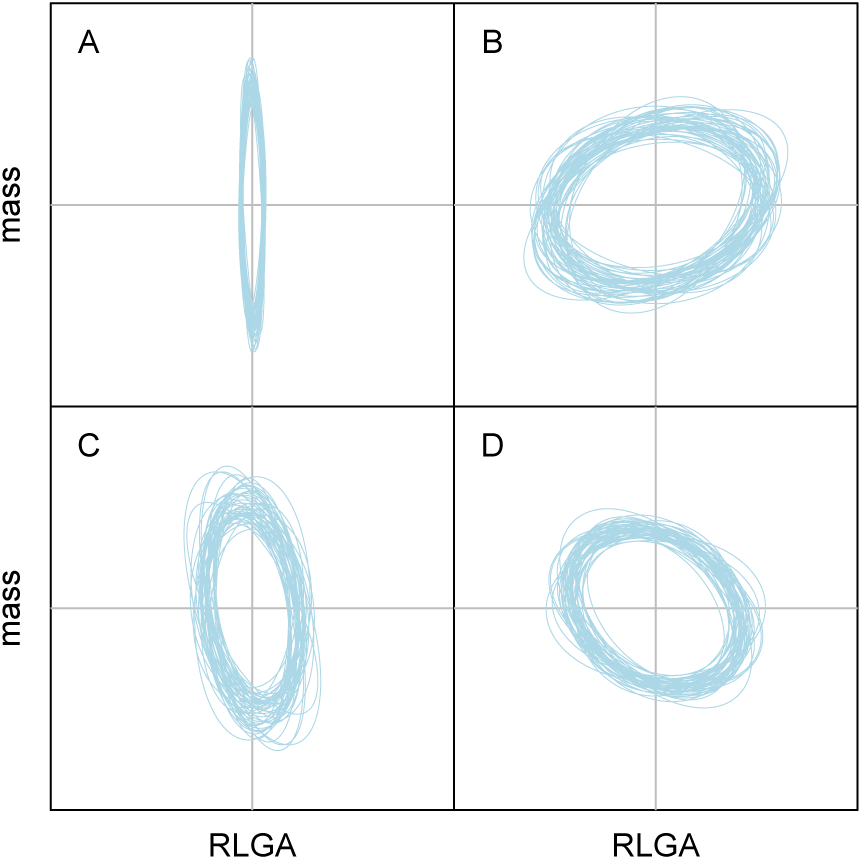
The 2×2 marginal distributions of **R**, restricted to body mass and relative lower grinding area (RLGA), can be obtained from the posterior distributions of the full 5×5 R matrices. Shown here are ellipses representing the 95% quantiles of 50 matrices sampled at random from the joint posterior distributions of A) cat–like hypercarnivores, B) weasel–like hypercarnivroes, C) omnivores, and D) mesocarnivores.

**Table 3:** Pairwise comparisons between the 2−2 **R** matrices for carnivorans belonging to four dietary regimes. P(Ψ*d*(A,B) *>* 0) is the probability that the two matrices define identical multivariate normal distributions. PP(rSDE) gives the posterior probability that matrix A is more eccentric (has a higher rSDE) than matrix B. ∠*AB* gives the mean angle between the principal eigenvectors of the two matrices; P(Ψ∠(A,B)*>* 0) is an associated test of whether they differ significantly. PP(disp) provides the posterior probability that the rate of multivariate dispersion is greater for regime A than for B. In each case, A refers to the first regime given and B refers to the second regime. Posterior probabilities are provided in A*>* B format for ease of presentation - it should be noted that PP *<* 0.05 is strong evidence for B *>* A.

| | $P(\Psi d(A,B) > 0)$ | $PP(rSDE)$ | $\angle AB$ | $P(\Psi \angle(A,B) > 0)$ | $PP(disp)$ | $PP(rlga)$ | $PP(mass)$ |
| --- | --- | --- | --- | --- | --- | --- | --- |
| Regime 1 vs 2 | <0.001 | 1 | 76 | <0.001 | 0.16 | 0 | 0.99 |
| Regime 1 vs 3 | <0.001 | 1 | 7 | 0.091 | 0.4 | 0 | 0.61 |
| Regime 1 vs anc | <0.001 | 1 | 53 | 0.001 | 0.39 | 0 | 0.99 |
| Regime 2 vs 3 | <0.001 | 0.03 | 68 | 0.003 | 0.78 | 1 | 0.04 |
| Regime 2 vs anc | 0.09 | 0.69 | 22 | 0.272 | 0.83 | 1 | 0.45 |
| Regime 3 vs anc | <0.001 | 0.99 | 45 | 0.016 | 0.5 | 0 | 0.98 |

This interpretation is further borne out by examination of evolutionary rates. Most inter-regime comparisons suggest little variation in rates of bivariate dispersion. We examined this lack of variation further by looking for differences between rate posterior distributions for RLGA and 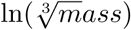 individually. Here, we found strong support (PP =1) for faster rates of relative lower grinding area evolution in omnivores and mesocarnivores compared with both hypercarnivorous regimes. For body mass, rates in the two hypercarnivore regimes were substantially faster than the omivores (PP[1 *>* 2] = 1, PP[3 *>* 2] = 0.96) and the mesocarnivores (PP[1 *> anc*]1, PP[3 *> anc*] = 0.97) while comparisons between hypercarnivores (PP1 *>* 3] = 0.76) and between mesocarnivores and omnivores (PP[*anc >* 2] = 0.53) reveal no significant differences.

## Discussion

### The Carnivoran General Adaptive Radiation

The order Carnivora underwent an early–burst of ecomorphological evolution, consistent with an Osbornian general adaptive radiation, but our ability to determine this is dependent on the traits studied. Of the individual ecomorphological traits we examined, only dental traits related to the relative proportion of non-meat items in the diet (Van Valkenburgh, 1988, 1991) showed signals consistent with an early burst model. The carnivoran carnassial has long been recognized as a key innovation that facilitated the dietary diversification of the clade from the Eocene onward (Matthew, 1909; Gregory and Hellman, 1939; Ewer, 1973) but it is interesting to note that functional traits related to shearing (P4P and RBL) accrued relatively modest support (*w_A_ <* 0.7) for the EB model compared to those related to grinding (*w_A_ >* 0.9). Studies of feeding behavior in wild carnivorans indicate that the carnassial blade is used primarily in cutting and chewing skin, and therefore plays an important role in allowing access to muscle, viscera, and bones within carcasses (Van Valkenburgh, 1996). Weak support for early burst dynamics in relative blade length and shape may therefore reflect the functional importance of a well-developed blade not only in hypercarnivorous taxa, but also in any taxon that seasonally hunts or scavenges vertebrate prey. The stronger support for EB dynamics in traits associated with the ability to process non-meat foods suggests that carnivorans represent an adaptive radiation along an axis of increasing dietary diversity that includes not only hypercarnivores, but also mesocarnivores, omnivores, and frugivores.

This dietary adaptive radiation characterizes the order as a whole however and its signal is not present in individual carnivoran families. Foote (1996) noted that such a pattern may occur where evolutionary rates have slowed sufficiently by time subclades originate that a signature of their decline is undetectable in trait data. Here though, failure to detect the carnivoran-wide early burst at lower levels seems to arise from variation in the mode of grinding area evolution within subclades, which results in dramatically different patterns of trait variation to those expected from the early burst inferred for the parent clade. For example, grinding area evolution in canids is characterized by a best fitting SSP model. Support for a model indicating low phylogenetic signal is consistent with repeated convergence on a constrained set of ecomorphological outcomes for this clade (Van Valkenburgh, 1991; Slater, 2015a), but is inconsistent with early burst dynamics. Best fitting Brownian motion models for malagasy carnivorans (Eupleridae) and mongooses (Herpestidae), in contrast, indicate that these clades have continued to expand in grinding area diversity at a slow but steady pace. The presence of such different macroevolutionary signatures at different levels of phylogeny suggests that early burst–like patterns may be an emergent property of complex, heterogeneous evolutionary dynamics, rather than resulting from simple slow–downs in evolutionary rate through time.

Such a perspective is logically consistent with conceptual models of early divergence into distinct adaptive zones combined with later diversification in and among sub-zones (Simpson, 1953; Valentine, 1980) but, if pervasive, has important implications for the validity of some phylogenetic tests of the early burst hypothesis. For example, the lack of temporal trends in the magnitude of per-branch rates of trait evolution has been offered as evidence that early bursts are an uncommon mode of adaptive radiation (Venditti et al., 2011). However, if evolutionary mode varies over a given phylogeny, then trends derived from per-branch rates estimated under a Brownian motion model may be extremely misleading. This is particularly true where equilibrial dynamics, such as Ornstein–Uhlenbeck or diversity–dependent processes, are involved, as the magnitude of trait change per unit time is not expected to be constant (Rabosky, 2009; Hunt, 2012). In our study, per-branch rates of grinding area evolution have indeed been quite variable through time and, contrary to expectations under early burst dynamics, the fastest rates occur in some of the youngest lineages such as *Lycalopex* canids (Figure 7A). However, branch durations also show a tendency to decrease towards the present (branch length = 1.26 + 0.33 parent node age, *R*^2^ = 0.385*, p <* 0.001), which will tend to result in larger per-branch rate estimates in young branches when the true mode of phenotypic evolution deviates from a constant–rates process. This phenomenon is analogous the the “Sadler effect” from sedimentary geology, where a trend of faster rates of sediment accumulation towards the present results from the discontinuous nature of sediment deposition combined with an increase in sequence length with increasing age (Sadler, 1981), and suggests that examining trends in per-branch rates might provide a poor test for early burst dynamics. More generally, it is worth stressing that a few recent, rapid rates do not negate a general partitioning of morphological variation among clades (Slater and Pennell, 2014; Cooney et al., 2017) which is the phenomenon we are actually interested in when studying general adaptive radiation. Relative subclade disparity through time (Harmon et al., 2003), though high for the first 20 Myr of carnivoran evolution where branch lengths are longest, gives way to a partitioning of variation among clades from 30 Ma until the present, coincident with the radiation of extant carnivoran families (Figure 7B). Integration of now extinct clades such as nimravids and amphicyonids, which diverged during the early phase of carnivoran diversification (Wesley-Hunt and Flynn, 2005), could increase the the signal for an early partitioning of disparity by breaking up long, old branches (Slater et al., 2012). Constant rates of grinding area evolution combined with highly selective, clade–specific patterns of extinction could plausibly generate this pattern, but it is difficult to contrive a realistic scenario in which the partitioning of variance among clades is overturned.

**Figure 7:**
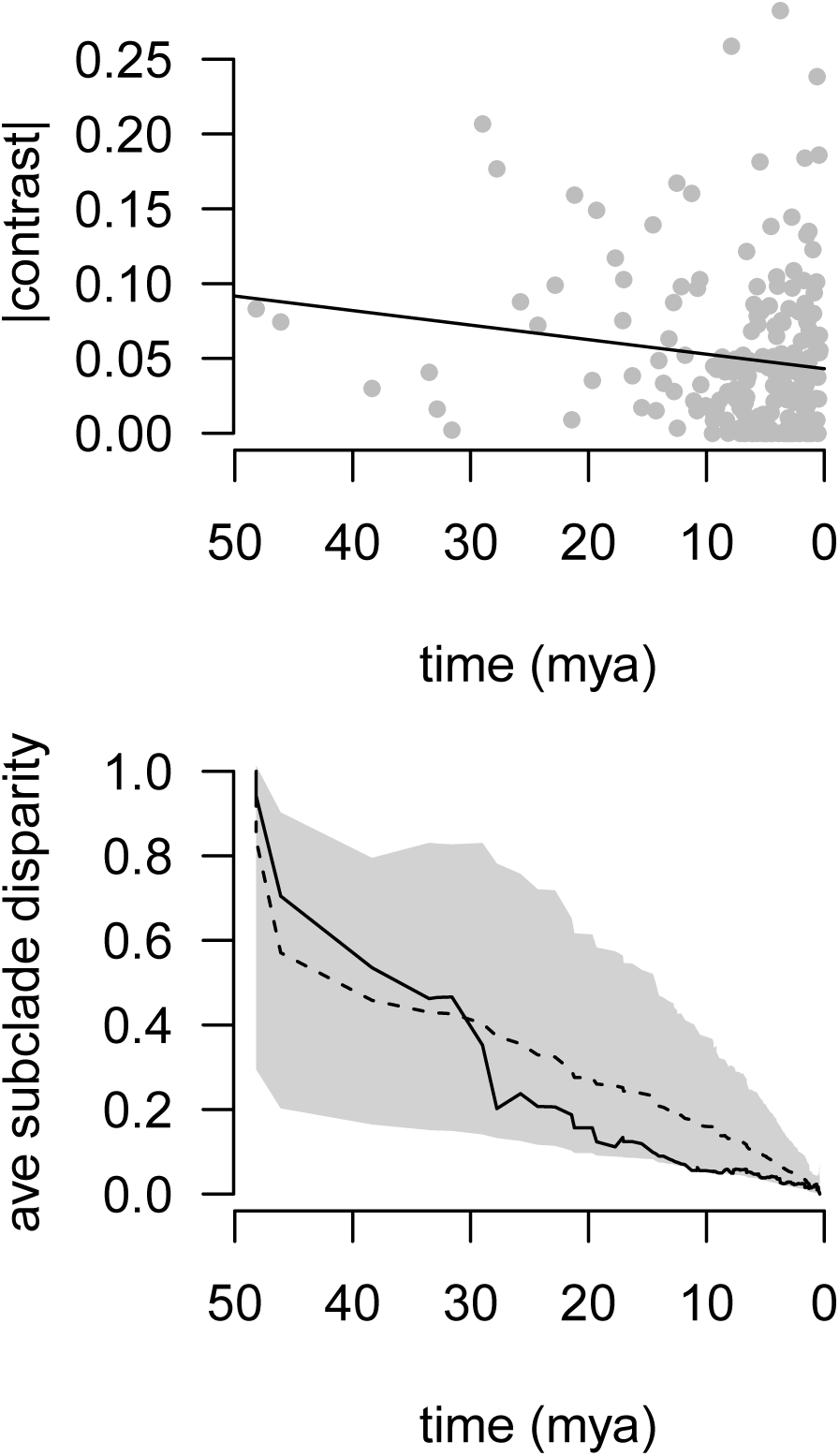
A) Rates of grinding area evolution slow down in general over carnivoran phylogeny but the fastest rates occur close to the tips. B) Despite fast rates along young branches, morphological disparity is increasingly partitioned among clades as one moves from root to tip, with a substantial drop at approximately 30 mya coincident with the origins of Feliformia and the onset of familial diversification.

### Body Size as an Axis of Local Adaptive Radiation

The lack of an early burst in carnivoran body mass evolution contrasts strongly with the signal detected in dental traits and corroborates the view that size can play an unpredictable role in macroevolutionary pattern (Jablonski, 1996). The use of size in macroevolutionary studies is often justified by the observation that many ecological, biomechanical and physiological traits show predictable patterns of allometric scaling (Bonner, 1965; Calder, 1984; Peters, 1986; Schmidt-Nielsen, 1984), and this is true in general terms for carnivorans where both prey size (Carbone et al., 1999, 2007; Van Valkenburgh, 1996) and the outcomes of interspecific interactions (Donadio and Buskirk, 2006) show strong size-based relationships. However, the weak correlation between mass and grinding area that emerges from our multivariate model fitting (model-averaged *r* = −0.086, Figure 5) corroborates a general decoupling of size and ecomorphology in Carnivora (Meloro and Raia, 2010) and implies that while size may play an important role over microevolutionary time scales, functional traits are more likely to carry a faithful signal of ecological diversification in the geologic past (Slater, 2015a). As a consequence, the lack of an early burst of body mass evolution in other clades should not be held as evidence against this mode of general adaptive radiation.

We find some support for the notion that body mass plays an important role in other forms of adaptive radiation. Osborn (1902) hypothesized that ongoing competition for resources within the lineages of a general adaptive radiation should result in local adaptive radiations, the evolution of assemblages of ecologically similar taxa with ecomorphological characteristics that facilitate resource partitioning. Our analyses suggest that body mass may be a particularly critical axis of local adaptive radiation in hypercarnivorous carnivorans, where the reduction or loss of molar grinding area reduces ecomorphological flexibility (Holliday and Steppan, 2004; Van Valkenburgh, 2007; Van Valkenburgh et al., 2004) and renders body mass the major axis of variation available to selection (Figure 6, Table 3). Indeed, extant and fossil hypercarnivore communities have long been recognized to exhibit strong size-based partitioning of prey resources and represent some of the few cases in which the presence of Hutchinsonian size ratios have been statistically established (Hutchinson, 1959; Dayan et al., 1990; Dayan and Simberloff, 1994; Werdelin, 1996; Meiri et al., 2005). Felid communities in both the Paleo- and Neotropics superficially appear to violate this prediction by consistently containing pairs of similarly sized taxa (Kiltie, 1984, 1988), though ecological data suggest that these pairs may avoid direct competition by spatio-temporally partitioning the environment (Bianchi et al., 2011), providing tantalizing cases of exceptions that prove the rule. It is important to stress, however, that despite the ecological importance of size in these clades and their faster rates of body mass evolution, there is no reason to expect strong phylogenetic signal or early burst-like evolutionary dynamics in traits underlying local adaptive radiations. Ecological mechanisms such as species sorting and character displacement are likely more important in generating equal size spacing (Rosenzweig, 1966; Davies et al., 2007; Pfennig and Pfennig, 2012) and we find no support for an early burst of body mass evolution in felids (*w_A_* = 0.19) or mustelids (*w_A_* = 0.2) when these clades are analyzed independently. Size may also provide a source of ecologically important morphological variation through allometry and some authors have suggested that size can, in this way, act as a line of least macroevolutionary resistance (Schluter, 1996a; Marroig and Cheverud, 2005, 2010). Birds provide a rare example of a clade with a particular propensity towards early bursts of body size evolution at both higher and lower phylogenetic levels (Richman and Price, 1992; Harmon et al., 2010; Jønsson et al., 2012) but some evidence suggests that allometric variation in cranial and beak morphology provides an important source of ecologically important variation (Bright et al., 2016), Thus, where early bursts of size evolution occur, they may represent instances of trait hitch-hiking (Jablonski, 2008, 2017), rather than divergence into distinct, size-based adaptive zones.

### Dimensionality of Adaptive Radiation

Although we find strong support for the early burst model of general adaptive radiation in univariate dental traits associated with dietary resource use, we find no support for early burst–like dynamics in multivariate ecomorphology. This result is surprising given that most paleobiological evidence for early burst–like dynamics comes from analyses of multivariate character data that characterize overall organismal form (Westoll, 1949; Foote, 1995; Hughes et al., 2013). Although several authors have shown that cladistic and morphometric data can provide similar characterizations of clade disparity (Villier and Eble, 2004; Anderson and Friedman, 2012; Foth et al., 2012; Hetherington et al., 2015) few have examined whether the choice of data affects inference of macroevolutionary mode. In a rare exception, Mongiardino-Koch et al. (2017) found conflicting signals when comparing cladistic and morphometric data for the scorpion genus *Brachistosternus* and interpreted the “late peak disparity” signal in their morphometric data to represent species–specific but convergent adaptations that overwrote historical patterns of ecomorphological evolution, such as early burst adaptive radiation. Support for the low phylogenetic signal, single stationary peak model for our multivariate ecomorphological data is consistent with this interpretation and helps reconcile our results with a disparity–based analysis of the North American carnivoran fossil record that showed no evidence for an early peak in multivariate ecomorphological variation (Wesley-Hunt, 2005). But, given the recent emergence and interest in multivariate phylogenetic methods for studying ecomorphological evolutoin (e.g., Clavel et al., 2015; Bastide et al., 2018), it seems reasonable to question why this distinction emerges and what it says about the dimensionality of different forms of adaptive radiation?

The ecological opportunity model of adaptive radiation is rooted in the idea that lineages within diversifying clades evolve rapidly towards novel or vacated peaks on the adaptive landscape (Schluter, 2000; Losos and Mahler, 2010). Conceptualizing this model in univariate space is straightforward as we can imagine isolated peaks or adaptive zones (Simpson, 1944, 1953) that directly relate morphology to function. Indeed, this idea is implicit in modern phylogenetic approaches for modeling adaptive evolution under an Ornstein-Uhlenbeck process (Hansen, 1997; Butler and King, 2004; Beaulieu et al., 2012) and is the pattern we recover here for relative grinding area. However, simulations show that where selection acts to optimize multiple functions simultaneously, the number of peaks on the multi-dimensional adaptive landscape, that is the number of phenotypic outcomes yielding comparable net fitness, increases while the fitness difference between the peaks and the valleys connecting them may decline. The result is a network of somewhat interconnected peaks that can easily be traversed over macroevolutionary timescales (Niklas, 1997, 1999). Marshall (2003, 2014) termed this outcome the “Principle of Frustration” because evolution’s ability to maximize all functions simultaneously is frustrated by trade–offs that result from the underlying morphological architecture of the organism. This idea is related to the concept of many-to-one mapping of form to function (Wainwright et al., 2005) where a massive amount of morphological variation may exist in individual components of an integrated functional system without affecting the functional output of the system itself. Critically, mosaicism in macroevolutionary rates and modes among different traits or body parts, a rather common pattern in fossil time series (Hopkins and Lidgard, 2012), is an expected consequence in such scenarios (Collar and Wainwright, 2006; Muñoz et al., 2017), and is the outcome we find here when comparing among best-fit models for individua ecomorphologicall traits. However, the weak evolutionary covariances among traits that result from such process will yield “noisy” multivariate data, where considerable morphological divergence is expected between young species pairs undergoing local adaptive radiation (Doebeli and Ispolatov, 2017). The choice of trait(s) is therefore critical when attempting to understand general, law–like behaviors in patterns of morphological evolution. Comparative evolutionary biologists must remember that all ecomorphological traits are not equal and those, like relative grinding area, that play a greater role in determining the fundamental ecological role of the organisms under study are more likely to be the ones in which early bursts of general adaptive radiation occur. Despite the greater potential to describe overall organismal form, multivariate data will not necessarily help detect general adaptive radiations and may even hinder the search for them.

## Conclusion

Phylogenetic comparative methods provide a powerful statistical framework for testing hypotheses about tempo and mode in phenotypic evolution, but the insights about how evolution works that can be gained from them depend wholly on the selection of appropriate data for the question at hand. Much of what we have learned about the prevalence of early bursts over the past 10 years has, we argue, been based on a conflating of processes occurring at different levels of the phylogenetic hierarchy and, as a result, we have made little progress in advancing our understanding of general law-like behaviors in morphological diversification (Gould, 1980). We here demonstrated that early burst dynamics characterize the evolution of a suite of traits associated with diet in the order Carnivora, but that this macroevolutionary signature characterizes the order and not individual families. Other traits, such as body mass, play critical roles in facilitating resource partitioning at lower phylogenetic levels but the evolutionary and ecological processes involved do not lead to early burst–like patterns in comparative data. Though challenges exist for assembling appropriate datasets (Cooney et al., 2017), restricting the expectation of an early burst of phenotypic evolution to Osbornian general adaptive radiations and identifying the traits involved and hierarchical levels at which this process operates (Erwin, 2000; Jablonski, 2007) offers a compelling route towards a more comprehensive understanding of the nomothetic forces shaping macroevolutionary pattern and process in morphological data.

## Acknowledgements

We thank curators at the Academy of Natural Sciences, Philadelphia, American Museum of Natural History, New York, Donald R Dickey collection, UCLA, Field Museum, Chicago, Los Angeles County Museum of Natural HistoryMuśeum national d’Histoire naturelle, Paris, National Museum of Natural History, Washington DC, Natural History Museum, London, and World Museum, Liverpool for access to specimens in their care. David Jablonski and Shauna Price provided valuable comments and criticisms of an earlier version of this manuscript.

## Data Archival Location

Phylogenetic and ecomorphological data and R scripts are deposited on DataDryad and will be made publically available on acceptance of the manuscript.

